# Phenotypic plasticity of antibiotic resistance, metabolism byproduct utilization and the evolution of mutually beneficial cooperation in *Escherichia coli*

**DOI:** 10.1101/2023.08.23.554550

**Authors:** Nan Ye, Beibei Hou, Jianxiao Song, Derek W Dunn, Zhanshan (Sam) Ma, Rui-Wu Wang

**Author notes:** For All Correspondence. Authors contributed equally (Nan Ye, Beibei Hou, Jianxiao Song).

## Abstract

Although tag-based donation and recognition have well explained how the cooperative individuals are positively assorted if the cooperative individuals possess some signals and are also able to detect such signals, an additional mechanism is required to explain why some individuals pay the costs of evolving such a tag that may not be rewarded subsequently, and how such tag-based cooperative individuals will meet other similar individuals with a very low mutation rate. Here, we show that many and even all *Escherichia coli* bacteria cells in the increased antibiotic concentration will plastically evolve to be antibiotic resistant individuals who could protect antibiotic sensitive strain from the attack of antibiotics, and the antibiotic resistant strain could reversibly evolve to be antibiotic sensitive in non-antibiotic supplement medium but in a harsher environment with low glucose. A further experiment showed that antibiotic-sensitive *E. coli* strain could in turn help reduce the concentration of indole produced by the resistant strain. This metabolic product is harmful to the growth of the antibiotic-resistant strain but benefits the antibiotic-sensitive strain by helping turn on the multi-drug exporter to discharge the antibiotic. The utilization of metabolism byproduct indole produced by antibiotic-resistant cells benefits antibiotic-sensitive cells, while the indole-absorbing service of antibiotic sensitive cells unconsciously help in nullifying the indole side effect on antibiotic resistant strain, and a mutual benefit cooperation could therefore evolve.

## Introduction

Kin selection and reciprocity are two well-known mechanisms for explaining the evolution of cooperation (Hamilton 1964, Maynard-Smith 1964, Trivers 1971). However, it is still unclear how cooperation evolves and persists in populations that are theoretically vulnerable to invasion by ‘cheaters’ (Smith & Schuster 2019, West et al., 2021), unrelated individuals who receive benefits from cooperative individuals but fail to reciprocate (Michod 1982, West et al., 2007). Tag-based donation and recognition theories posit that cooperative individuals may become associated with each other and thus form groups via positive assortment through a tag such as a recognizable phenotypic trait (Dawkins 1976, Jansen & Van Baalen 2006, Kim 2010, Laird 2011, West et al., 2021). That cooperative individuals easily meet and/or associate with one another is a key mechanism for positive assortment of cooperative individuals (Pepper & Smuts 2002, Joshi et al., 2017), especially in large populations. Tag based theories, however, need additional mechanisms to explain how a mutant with a specific tag is initially selected and then meets other similar mutants, especially when the mutation rate is low (Lynch et al., 2016).

The bacterium *Escherichia coli* (MG 1655) is a widely-used model organism to study how individual cells cooperate within a population (Lee et al., 2010). The wild-type *E. coli* (MG1655) is sensitive to antibiotics and its growth is much constrained by exposure to norfloxacin via both nucleic acid replication and transcription (Baquero & Levin 2021). However, if *E. coli* MG1655 is incubated in an antibiotic-culture medium, resistant cells (*E. coli*-R) will be selected and further benefit the wild type cells (antibiotic sensitive strain, *E. coli*-S) through the production of indole, which could increase bacterial antibiotic resistance by turning on the efflux pumps and oxidative-stress protective mechanisms (Lee et al., 2010).

Benefits accrued by wild-type *E. coli* from antibiotic inhibition by antibiotic-resistant *E. coli* is often regarded as an example of altruism (Lee et al., 2010, Carmona-Fontaine & Xavier 2012, Cabral et al., 2018). However, how antibiotic-resistant cells respond to selection and become positively assorted in a population remains poorly understood. Here, we show that some wild-type cells in *E. coli* population evolved into ‘altruistic’ individuals via phenotypic plasticity under antibiotic pressure, and the process is rapid and reversible. Our experiments also showed a reciprocal exchange between the metabolism byproduct indole of antibiotic resistant cells and the indole-absorbing service of antibiotic sensitive cells. Each of these two modes of cooperation are unconscious cost-free provisions to partners during the early stage of cooperation. Such positive feedback will likely facilitate the evolution of the potentially costly reciprocal behavior, which has been previously demonstrated in both *E. coli* (Preussger et al., 2020) and *Salmonella enterica* (Harcombe et al., 2018).

## Materials and methods

### 1. Selection of norfloxacin resistant *E. coli*

A ‘wild-type’ antibiotic-sensitive *E. coli* MG1655 strain (*E. coli*-S) was obtained from the National Center for Type Culture Collection (CCTCC), and incubated in a lysogeny broth liquid medium (LB) supplemented with different concentrations of norfloxacin (details below). The antimicrobial susceptibility of this *E. coli*-S strain was determined by initial streaking on norfloxacin gradient plates, followed by incubation for 15 h at 37 L. The tested tolerance threshold of this *E. coli*-S strain to norfloxacin was 150 ng mL^−1^. The *E. coli*-S culture was inoculated to an LB plate with 150 ng mL^−1^ norfloxacin for 24 hours (at 37 L). A single colony was then inoculated to an LB plate with 300 ng mL^−1^ norfloxacin. By continuously culturing with increasing concentrations of norfloxacin, the resulting resistant strain was able to survive on LB medium with 2400 ng mL^−1^ norfloxacin. After more than three times isolation and purification, this antibiotic-resistant strain (*E. coli*-R) was stored in 25% glycerol at −80 L for future experimentation.

### 2. Effect of different norfloxacin concentrations on the proportion of norfloxacin tolerant bacterial cells

20 µl (OD_600_ ∼ 0.01) of *E. coli*-S was inoculated to 5 mL falcon tubes with LB liquid mediums at a norfloxacin concentration of either 0, 37.5, 75, and 150 ng mL^−1^ (four replicates per treatment). Following incubation for 12 hours at 37 L, OD_600_ (optical density at 600nm) of each replicate was measured and diluted to the same value. 50 µl of diluted culture was then transferred to fresh LB liquid medium with norfloxacin concentration at 0 or 600 ng mL^−1^ (four replicates per treatment). The proportion of norfloxacin-resistant (*E. coli*-R) cells was calculated as OD_600_ (norfloxacin=600 ng mL^−1^)/ OD_600_(norfloxacin = 0 ng mL^−1^) in LB liquid culture (Supplementary Figure S1). The same experiment was also conducted using LB agar plates and the proportion of norfloxacin-resistant bacterial cells was measured by colony-forming units (CFUs) (Supplementary Figure S3).

### 3. Phenotypic plasticity assay

We further tested if the norfloxacin resistant *E. coli*-R could reversibly evolve to be antibiotic sensitive in a norfloxacin-free medium. Specifically, we transferred *E. coli*-R (OD_600_ ∼ 0.001) into two fresh media treatments for seven days (six replicates for each treatment). The first treatment was a low nutrition (0.05% glucose) M9 medium (6.8 g/L Na2HPO4, 3 g/L KH2PO4, 1 g/L NH4Cl, 0.5 g/L NaCl, 2 mM MgSO_4_·7H_2_O, 0.5 mM thiamine hydrochloride, 0.0025 g/L FeSO_4_·7H_2_O), and the second a full nutrition (0.5% glucose) M9 medium (otherwise identical to the low nutrition medium). In both treatments norfloxacin was absent.

After incubation at 37 ⃩, the proportion of *E. coli*-R was calculated as described above.

### 4. Growth *E. coli*-R and *E. coli*-S strains incubated either in isolation or together under variable norfloxacin conditions

*E. coli*-S labeled with the green fluorescent protein (gfp) gene and *E. coli*-R labeled with the red fluorescent protein (rfp) gene were inoculated to 5mL LB liquid medium followed by overnight incubation at 37°C. The two overnight cultures were diluted to the same value at OD_600_. Then, *E. coli*-S and *E. coli*-R were mixed at a ratio of 99:1 in a 96-well cell culture plate with 190 μl LB liquid medium and 2400 ng mL^−1^ norfloxacin (four replicates per treatment). Simultaneously, 9.9 μl *E. coli*-S or 0.1 μl *E. coli*-R were inoculated in a 96-well cell culture plate with 190.1 μl or 199.9 μl LB liquid medium, respectively, with the addition of 2400 ng mL^−1^ norfloxacin (four replicates per treatment). A blank control only contained 200 μl LB liquid medium. All replicates (N = 4) were cultured for 36h in a multifunctional microplate reader and the OD value was measured at hourly intervals (OD_600_).

### 5. Determining the composition of mixtures

*E. coli*-S with gfp and *E. coli*-R with rfp were inoculated to 5mL of LB liquid medium followed by overnight incubation at 37 L. The two overnight cultures were diluted to the same value at OD_600_. Then, *E. coli*-S and *E. coli*-R were mixed at a ratio of 99:1, and added to 5 mL of LB liquid medium supplemented with 1500 ng mL^−1^ norfloxacin followed by 17 h incubation at 37 L (three replicates per treatment). Simultaneously, 99μl *E. coli*-S or 1μl *E. coli*-R were inoculated separately to 5 mL of LB liquid medium supplemented with 1500 ng mL^−1^ norfloxacin, followed by 17h incubation at 37 L (three replicates). Each culture was diluted to 10^4^ cells per mL with PBS buffer at 9h, 11h, 13h, 15h, 17h. Flow cytometry (FACSAria III) was then used to calculate the total number of *E. coli*-S and *E. coli*-R cells. The laser and filter configurations for GFP and RFP with flow cytometer were 50 mW 488 nm laser with 505nm filter for GFP and 588 nm filter for RFP (Hart et al., 2019).

### 6. Extracellular indole quantification

Monocultures of *E. coli*-S and *E. coli*-R strains were each grown separately to OD_600_ ∼ 0.5 in LB medium at 37 °C and 220 rpm. Cell cultures (1 mL) were collected at 6, 12 and 18 h, centrifuged at 14,000 rpm for 60 minutes, and the supernatant collected for indole quantification. Extracellular indole was measured by reverse-phase HPLC using a C18 Waters Spherisorb ODS-2 column (25 cm × 4.6 mm, 3 µm) set at 40 °C and gradient elution with acetonitrile-0.1% (v/v), formic acid and H_2_O-0.1% (v/v) formic acid at the mobile phases at a flow rate of 1 mL min^−1^ (35:65 for 0-5 min, 65:35 for 5-12 min, 35:65 for 12-13 min, and 35:65 for 13-30 min). Under these conditions, the retention time for indole was 13.4 min, and the absorbance maximum was 271 nm.

### 7. Effects of indole on the growth of *E. coli*-S and *E. coli*-R in the presence of norfloxacin

Indole was dissolved in methanol to a concentration of 50 mmol L^−1^. Different norfloxacin concentrations (37.5, 75 and 150 ng mL^−1^) were added to 5 mL of LB liquid medium with or without the presence of 500 µmol L^−1^ indole. 50 μl of *E. coli*-S (OD_600_∼0.5) were inoculated to each treatment (three replicates per treatment). 50 μl of *E. coli*-R (OD_600_∼0.5) was also inoculated to 5mL LB liquid medium supplemented with norfloxacin at 1200 ng mL^−1^ or 2400 ng mL^−1^ (three replicates per treatment). For each norfloxacin treatment, 0 or 500 µmol L^−1^ indole was added. OD_600_ of each replicate was measured after 12 hours of incubation at 37 L and 220 rpm.

### 8. Statistical analysis

Shapiro-Wilk and Levene’s test were used to test for normality and homogeneity of variance, respectively. Parametric tests were used when error variances of the data were normally distributed. Otherwise, a non-parametric equivalent was used. The test used and sample size (n) are specified in the results or figure captions. Asterisks indicate significant differences in pairwise comparisons (*p < 0.05, **p< 0.01, ***p< 0.001). All analyses were conducted in SPSS 22.0 (SPSS Inc., Chicago).

## Results

### 1. Resistant phenotypes of *E. coli* as a result of phenotypic plasticity

In the norfloxacin selection experiment, we found that the higher initial norfloxacin concentration in culture, the higher proportion of *E. coli*-R cells in the total bacterial population were observed in 600 ng mL^−1^ norfloxacin after 12 hours (Fig. 1). After exposure to 150 ng mL^−1^ norfloxacin, almost all of the cells were able to tolerate 600 ng mL^−1^ norfloxacin (Fig. 1). Similar results were apparent when the experiment was repeated with ampicillin, kanamycin and tetracycline (Fig. S2&S3).

**Fig. 1.**
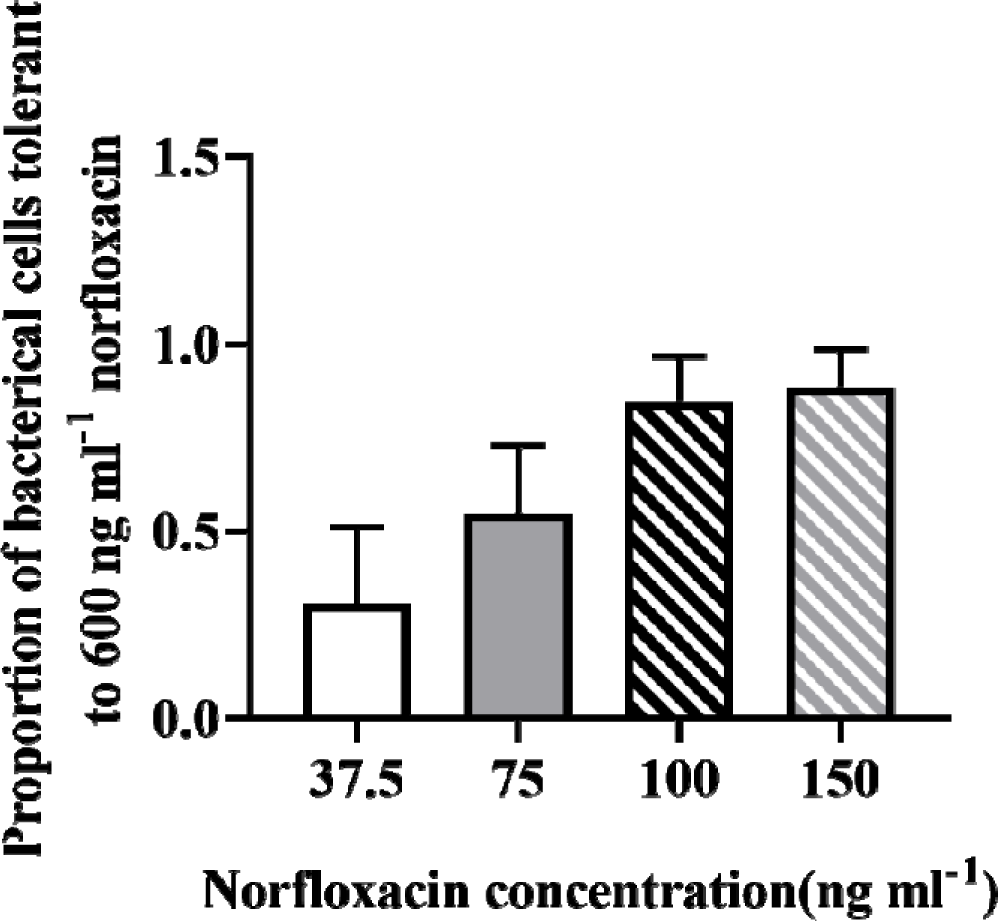
Proportion of bacterial cells tolerant to 600 ng mL^−1^ norfloxacin after 12 hours incubation in which the samples were exposed for 12 hours to either 37.5, 75, 100, or 150 ng mL^−1^ norfloxacin.

These *E. coli*-R cells, however, could phenotypically ‘reverse’ and shift to be antibiotic sensitive. In M9 medium without norfloxacin, some *E. coli*-R cells will not evolve to be antibiotics sensitive. However, the proportion of *E. coli*-R decreased from 90% to 2% in M9 medium without norfloxacin but with lower glucose (0.05%) than normal (Fig. 2), indicating that the majority of antibiotic-resistant cells lost their resistance in the nutrient-poor environment.

**Fig. 2.**
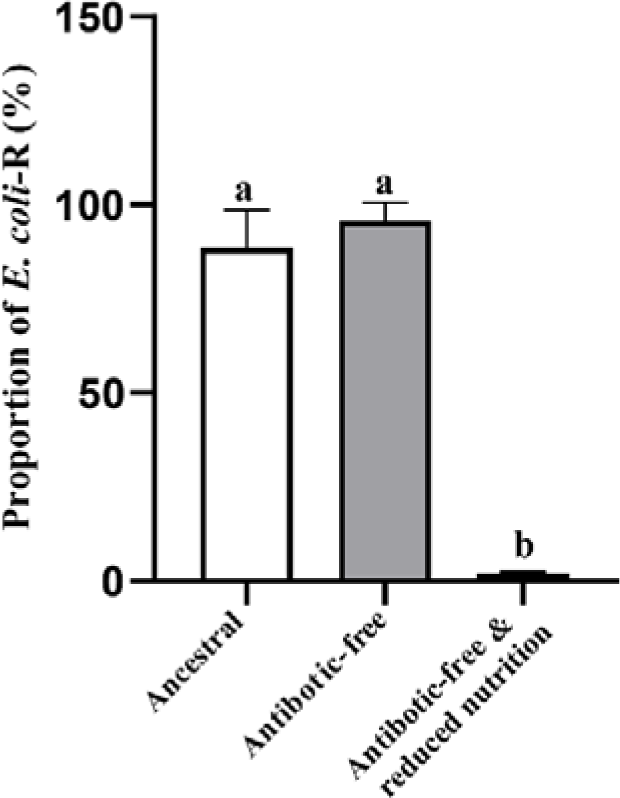
Reduced proportion of *E. coli*-R (%) after removal of norfloxacin. Ancestral: Resistant bacteria that can grow in 600 ng/ml norfloxacin after 150 ng/ml norfloxacin domestication in Figure 1. Antibiotic-free: *E. coli*-R incubated without norfloxacin for seven days; maintained high resistance after a few generations of growth in antibiotic-free conditions. Antibiotic-free and reduced nutrition: ancestral *E. coli*-R cells incubated in low nutrition (0.05% glucose) medium without norfloxacin. Mean ± SE (standard error) values are shown. Different letters indicate significant difference among groups (One way ANOVA, p < 0.001, F = 484.487, df = 15).

### 2. Mutual benefit between *E. coli*-S and *E. coli*-R in mixed population

We then tested the effect of antibiotics on both *E. coli*-S and *E. coli*-R in isolation or together. In a mixed population consisting of *E. coli*-S and *E. coli*-R beginning at a ratio of 99:1, respectively, the total biomass (measured as OD_600_) and growth patterns of the two-strain cultured population (*E. coli*-R and *E. coli*-S together) differed to those of each single-strain (*E. coli*-R or *E. coli*-S) population (Fig. 3). From 8 to 20 hours after population establishment (exponential phase), the population growth rate and total biomass of the single-strain *E. coli*-R culture was higher than the mixed, two-strain culture containing both *E. coli*-R and *E. coli*-S. After 20 hours (stationary phase) both mixed and *E. coli*-R only strain populations ceased to increase and reached an asymptote, with the biomass of the mixed strain exceeding that of the single *E. coli*-R only strain. When *E. coli*-S was cultured alone, total biomass remained unchanged throughout the duration of the experiment (36 hours), consistent with antibiotics effectively constraining population growth (Fig. 3).

**Fig. 3.**
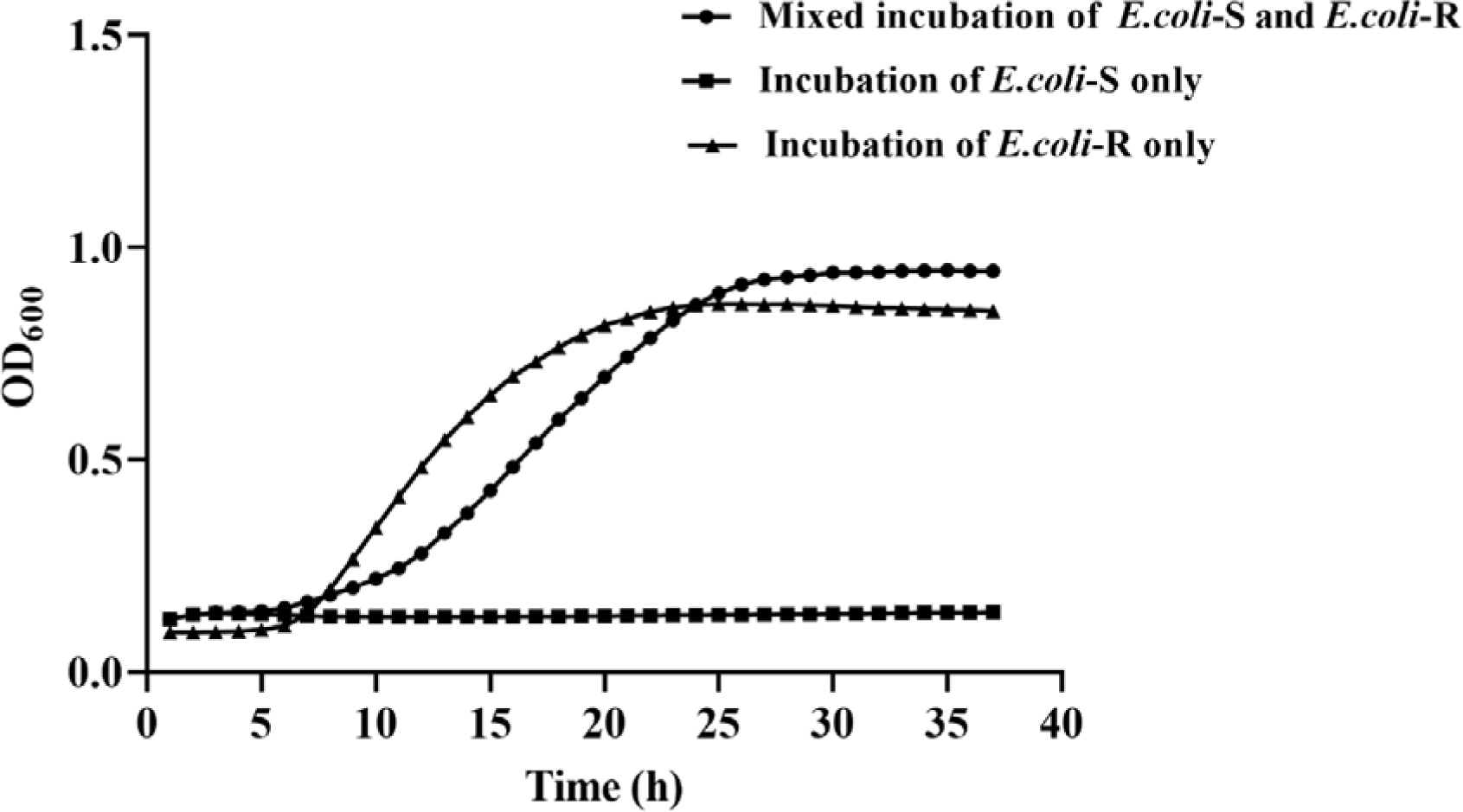
Growth curves of *E. coli* measured as OD_600_ in 2400 ng mL^−1^ norfloxacin culture. In mixed incubation, the initial ratio between of *E. coli*-S and *E. coli*-R is 99:1.

We then tested if the mixed cultures in an antibiotic medium could enhance the growth of *E. coli*-S and *E. coli*-R. To do so, we grew monocultures and co-cultures of *E. coli*-S and *E. coli*-R in LB medium supplemented with 1500 ng mL^−1^ norfloxacin. In the early growth stage (0-20 hrs.), *E.coli*-R cell numbers increased from 493±63 at 9 hours to 57034±15564 at 15 hours (average ± standard deviation, n=3); although slightly higher than those in the *E. coli*-R only population at 11, 13 and 15 hours, differences were not statistically significant (Fig. 4A, independent samples t-test, 11h : t = −1.048, p = 0.232,, n = 3; 13h: t = −9.11, p = 0.414, n = 3; 15h : t = −1.291, p = 0.290, n = 3). Cell numbers of *E. coli*-S were higher in the mixed-strain population than in the *E. coli*-S only population at all sampling times (Fig. 4B). These data suggest that the presence of antibiotic resistant *E. coli*-R enhanced survival and/or reproduction of *E. coli*-S under antibiotic stress.

**Fig.4.**
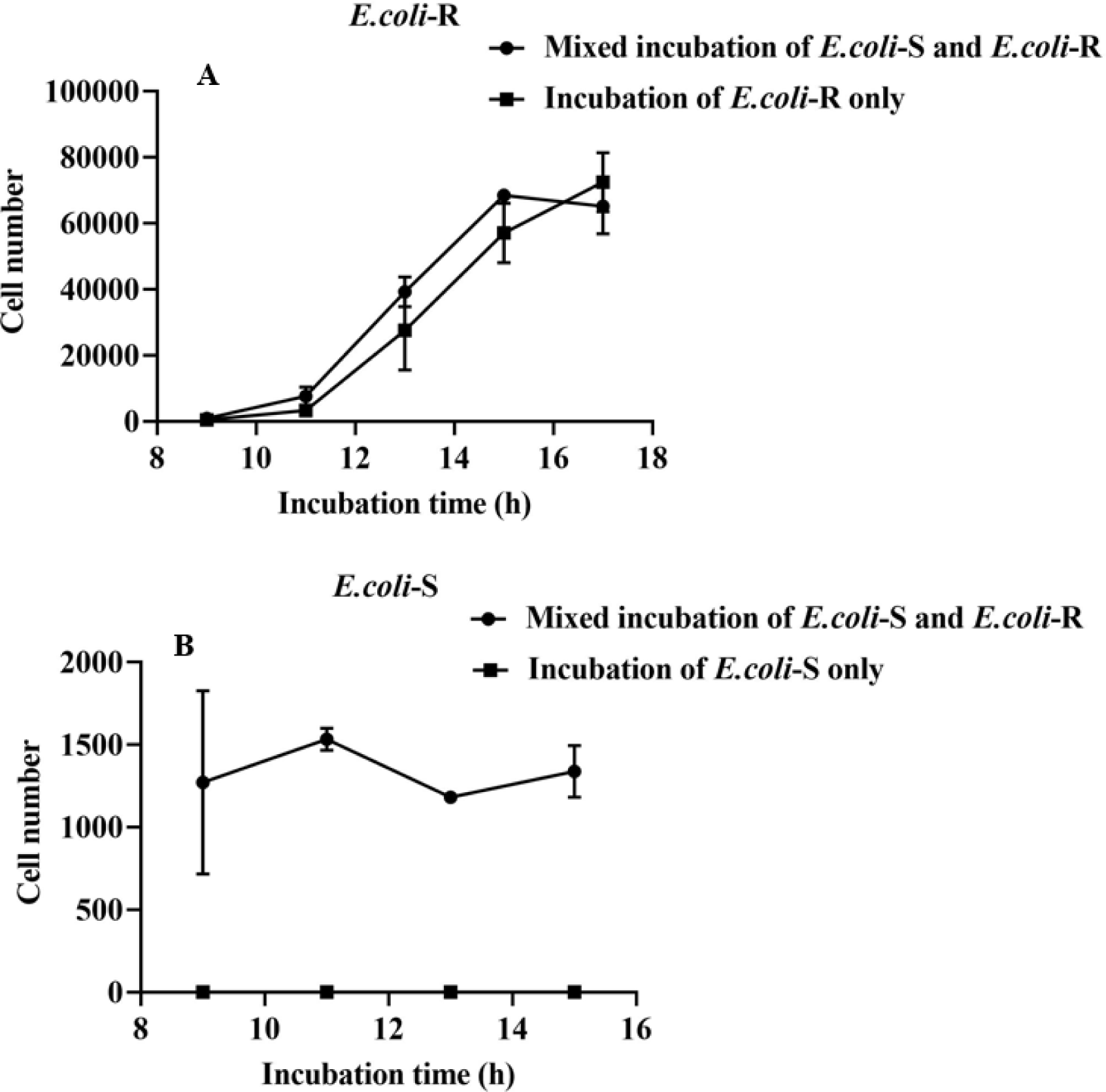
Cell numbers of both *E. coli*-R (A) and *E. coli*-S (B) counted by a flow cytometer. In mixed-strain populations, the initial ratio between *E. coli*-S and *E. coli*-R was 99:1. The norfloxacin concentration was 1500 ng mL^−1^ for all three populations.

We also found that the addition of indole inhibits population growth of *E. coli*-R but can promote growth of *E. coli*-S. In the presence of norfloxacin, indole concentration was 420 µmol L^−1^ in the mixed-strain culture, significantly lower (one-way ANOVA, n = 3, p = 0.001) than indole (450 µmol L^−1^) in the *E. coli*-R monoculture (Fig. 5). When 500 µmol L^−1^ indole was added into monocultures of *E. coli*-S or *E. coli*-R at different concentrations of norfloxacin, at low antibiotic concentrations (0, 37.5 ng mL^−1^ and 75 ng mL^−1^, Supplementary Figure S4, S5), indole did not significantly affect growth of *E. coli*-S (independent samples t-test comparing optical densities with 500 µmol L^−1^ indole and without indole during incubation, 37.5 ng mL^−1^: p = 0.021, t = −.674, n = 3; 75 ng mL^−1^: p = 0.256, t = 1.323, n = 3). However, when cultured with 150 ng mL^−1^ norfloxacin, the biomass of *E. coli*-S in the culture with 500 µmol L^−1^ indole was significantly higher than the culture in which indole was absent (independent samples t-test: p = 0.002, t = −7.431, n = 3, Fig. 6). This showed that the addition of indole, at high antibiotic concentrations, may result in increased *E. coli*-S biomass. However, additional indole significantly reduced *E. coli*-R total biomass compared with controls with only 1200 ng mL^−1^ or 2400 ng mL^−1^ norfloxacin (independent samples t-test, 1200 ng mL^−1^: p = 0.005, t = 5.590, n = 3; 2400 ng mL^−1^: p = 0.013, t = 4.243, n = 3). In the absence of norfloxacin, indole also significantly inhibited *E. coli*-R growth (Supplementary Figure S5B).

**Fig. 5.**
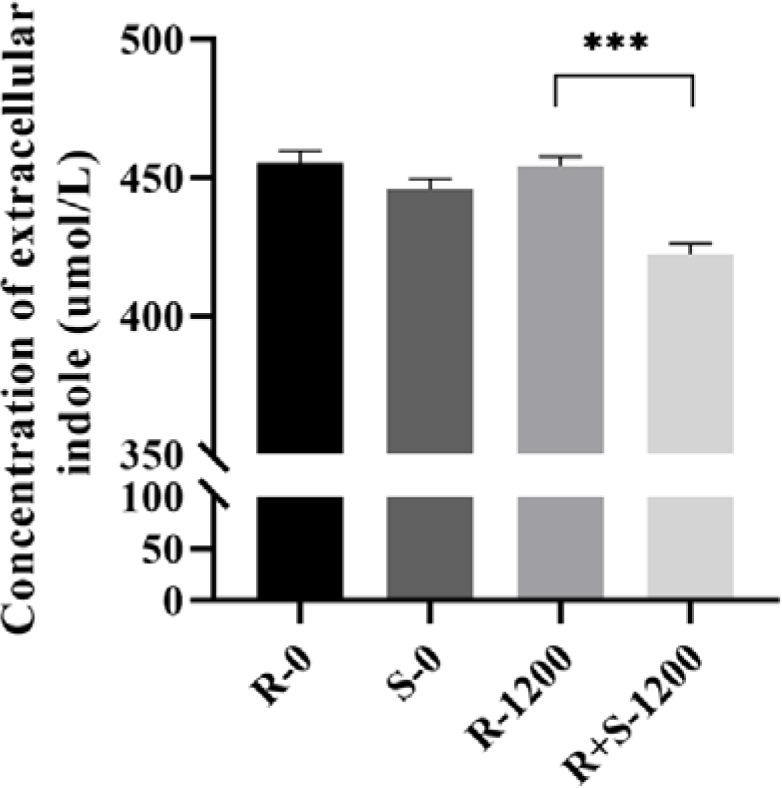
Concentration of extracellular indole in different treatments of *E. coli*. R-0: norfloxacin absent, *E. coli*-R. S-0: norfloxacin absent, *E. coli*-S. R-1200: 1200 ng mL^−1^ norfloxacin, *E. coli*-R. R+S-1200: 1200 ng mL^−1^ norfloxacin, *E. coli*-S and *E. coli*-R combined (99:1).

**Fig 6.**
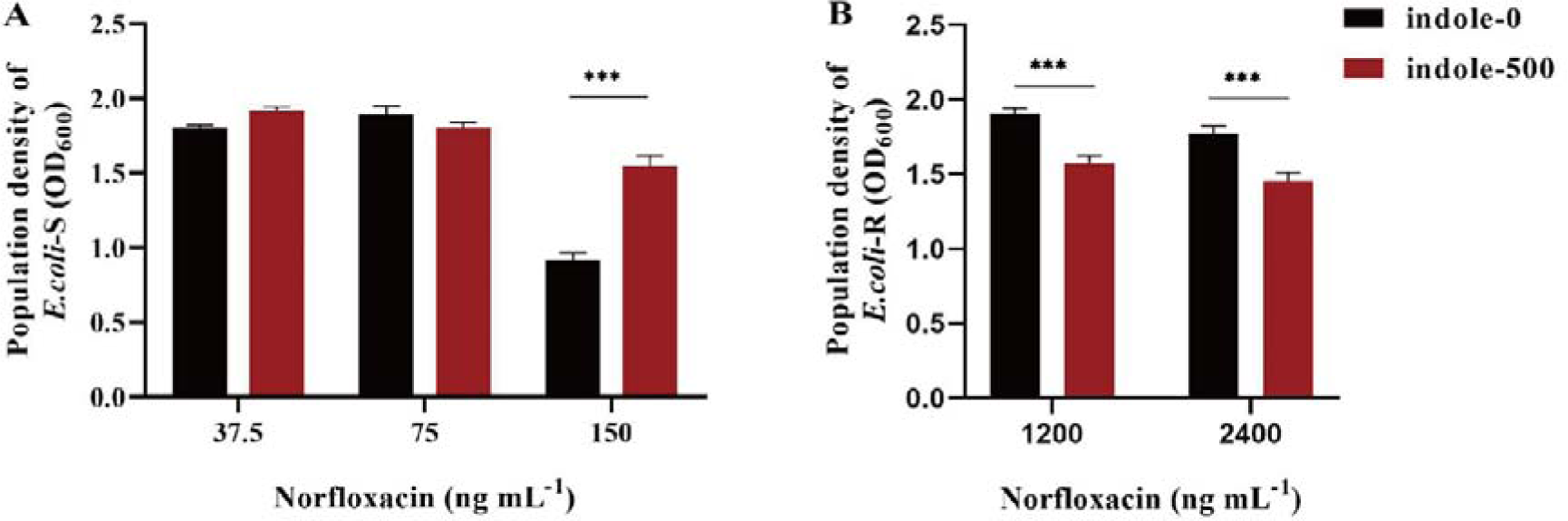
The effects of indole on the growth of *E. coli*-S (A) and *E. coli*-R (B) at different concentrations of norfloxacin. Indole was added to each treatment at a concentration of 500 µmol L^−1^. The norfloxacin concentrations were either 37.5, 75, 150 ng mL^−1^(A), and 1200 or 2400 ng mL^−1^ (B). All treatments were incubated in Luria-Bertani (LB) medium.

## Discussion

Antibiotic-resistant *E. coli* strains can survive under antibiotics due to either genetic resistance and or phenotypic tolerance (Wiuff et al., 2005, Jayaraman 2008, Munita & Arias 2016). Genetic resistance can be the result of natural selection favoring beneficial gene mutations, genes acquired via recombination, and gene acquisition mediated by transfer elements (e.g. via plasmids) (Thomas & Nielsen 2005, Didelot & Maiden 2010, Kivisaar 2020). However, our results showed that most of the *E. coli* cells exposed to high concentrations of antibiotic will quickly evolve to be antibiotic resistant individuals (Fig. 1), resistance likely to be a result of phenotypic resistance (tolerance). This is further confirmed by our finding that *E. coli*-R lose antibiotic-resistance at low glucose levels in the absence of antibiotics. Our data suggests that most of the *E. coli*-R cells are more likely to acquire antibiotic resistance through phenotypically plastic type (Wiuff et al., 2005), since genetic mutations usually generate non-reversible phenotypes (Sandoval-Motta & Aldana 2016). Such phenotypic plasticity may have resulted from increased cell membrane permeability under antibiotic stress (Wax et al., 2007).

Our results suggest that “altruistic” antibiotic-resistant cells might unconsciously benefit the antibiotic-sensitive cells through the metabolite byproduct indole, helping turn-on the multi-drug exporter of antibiotic-sensitive cells. The utilization of indole by antibiotic-sensitive *E. coli* may also reduce the indole overall concentration. Because indole is potentially harmful to antibiotic-resistant cells due to inhibiting cell growth / division in *E. coli*-R (Fig. 6B), antibiotic-resistant cells may accrue benefits from the actions of *E. coli*-S (Chimerel et al., 2012, Kim & Park 2015). Through such active indole absorption by *E. coli*-S, the accumulation of intracellular indole from *E. coli*-R will decrease, thus reducing the toxicity of indole to *E. coli*-R cells. Under antibiotic-stressed conditions, mutually beneficial cooperation between resistant and wild-type *E. coli* strains will thus result (Fig. 7). In the early stages of this cooperative interaction, there are no clear costs to cooperation for both *E. coli*-R and *E. coli*-S. However, *E. coli*-R might subsequently evolve over several generations to actively pay costs of benefiting wild-type individuals because of positive feedback from *E. coli*-S. Previous studies on the evolution of bacterial cooperation (*E. coli* and *Salmonella enterica*) have shown that byproduct consumption (a passive process) will eventually evolve into costly reciprocal cooperation (an active process) with increased positive fitness feedbacks among interacting species (Harcombe et al., 2018, Preussger et al., 2020).

**Fig. 7:**
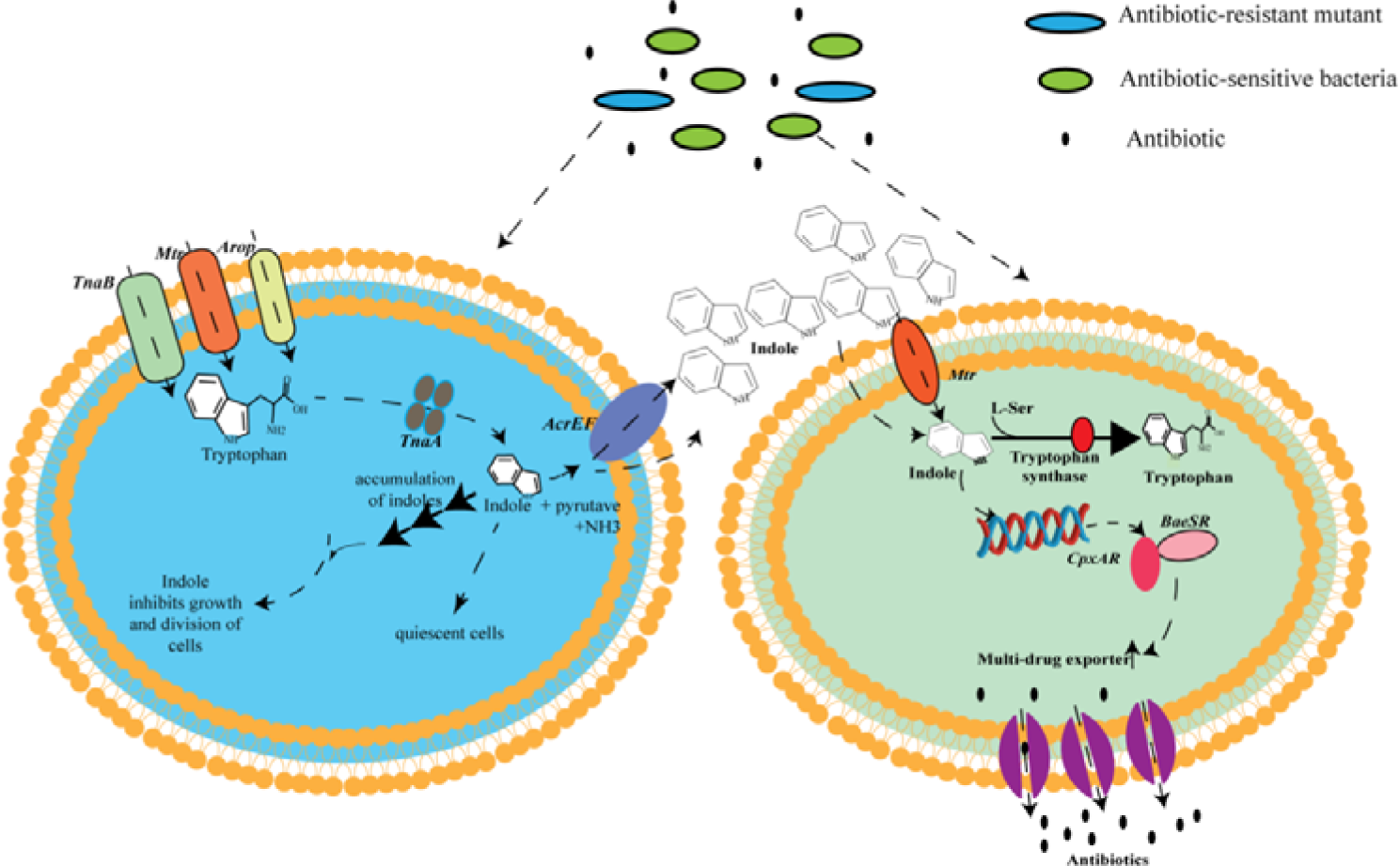
The mechanisms driving reciprocal cooperation between the antibiotic-resistant strain and the antibiotic-sensitive strain of *E. coli*. In an antibiotic-rich culture, *E. coli* R is still active in cell division and may therefore continue to synthesize indole from tryptophan via tryptophanase (*TnaA*). The production of indole by the *E. coli* R strain will diffuse across the cell membrane, with any accumulation of high concentrations of indole inhibiting growth and cell division of *E. coli* R. However, the production of indole by *E. coli* R may help *E. coli* S to initiate some transcription and translation processes to turn on the multi-drug exporter. Therefore, any consumption of *E. coli* S will reduce the concentration of indole in the wider environment thus benefiting *E. coli* R cell growth.

Phenotypic plasticity provides a new mechanism on how cooperative actors may become assorted over a relative short time period. In our *E. coli* experiments, the proportion of antibiotic-resistant cells increases according to antibiotic concentration, with all wild-type *E. coli* cells evolving to be antibiotic-resistant in a 150 ng mL^−1^ norfloxacin medium (Fig. 1).

The antibiotic-resistant *E. coli* cells may ‘behave’ differently (e.g., slow growing (Jayaraman 2008)) or have a different morphology (e, g., longer or wider flagellum; elongation of cells (Bos et al., 2015, Frost et al., 2018)) under antibiotic stress, which may act as a ‘tag’ to enable *E. coli* R to become positively assorted. Such behavior or morphological variation would thus be a side-effect of ‘cooperative’ antibiotic-resistant individuals at the macro level under antibiotic stress but would not involve any costs of benefitting the wild strain (macro level, cooperative).

The metabolite indole is the likely resource involved in this mutual beneficial cooperative interaction between *E. coli*-R and *E. coli*-S. As a metabolic byproduct of *E. coli* cell growth, indole may be still produced by the antibiotic resistant strain when exposed to antibiotics because antibiotic-resistant cells could still divide. Our data also show that both *E. coli*-R and *E. coli*-S are capable of producing up to approximately 500 µmol L^−1^ of indole in the absence of antibiotics, which is consistent with previous reports (Yanofsky et al., 1991). However, when stressed due to exposure to antibiotics, *E. coli*-S will not produce indole due to the suppression of cell division. *E. coli*-R will maintain indole production, even when exposed to high antibiotic concentrations. In mixed populations of *E. coli*-R and *E. coli*-S under antibiotic stress, indole produced by *E. coli*-R will enter *E. coli*-S cells via the cell membrane by diffusion or a channel protein (Walter et al., 2020, Zarkan et al., 2020). Within *E. coli*-S cells, indole will subsequently turn-on efflux pumps or initiate any oxidative-stress protective mechanisms (Lee et al., 2010, Kim & Park 2015, Munita & Arias 2016), which may enhance *E. coli*-S survival and/or reproduction (Fig. 6A).

## Acknowledgments

This research was supported by NSFC-Yunnan United fund (U2102221), the National Science Fund for Distinguished Young Scholars (31325005), the National Natural Science Foundation of China (31170408, 31270433, 31370408; 31360104; 32070453), the Innovation Foundation for Doctor Dissertation of Northwestern Polytechnical University (CX2023097). The authors thank Marco Archetti and Wenying Shou for critical comments on the manuscript.

## Authors contributions

RWW, JS designed the experiments, RWW, JS, NY, DWD wrote the manuscript, JS, NY, BH conduct the experiments and data analysis.

## Competing interests

The authors declare no competing interests.

## Notes

### Competing Interest Statement

The authors have declared no competing interest.

## References

Baquero, F., and B. R. Levin. 2021. "Proximate and ultimate causes of the bactericidal action of antibiotics." Nature Reviews Microbiology 19:123–132.

Bos, J., Q. Zhang, S. Vyawahare, E. Rogers, S. M. Rosenberg, and R. H. Austin. 2015. "Emergence of antibiotic resistance from multinucleated bacterial filaments." Proceedings of the National Academy of Sciences 112:178–183.

Cabral, D. J., J. I. Wurster, and P. Belenky. 2018. "Antibiotic Persistence as a Metabolic Adaptation: Stress, Metabolism, the Host, and New Directions." Pharmaceuticals 11:14.

Carmona-Fontaine, C., and J. B. Xavier. 2012. "Altruistic cell death and collective drug resistance." Molecular systems biology 8:627.

Chimerel, C., C. M. Field, S. Piñero-Fernandez, U. F. Keyser, and D. K. Summers. 2012. "Indole prevents Escherichia coli cell division by modulating membrane potential." Biochimica et Biophysica Acta (BBA)-Biomembranes 1818:1590–1594.

Dawkins, R. 1976. The selfish gene. Oxford University Press, New York.

Didelot, X., and M. C. Maiden. 2010. "Impact of recombination on bacterial evolution." Trends in microbiology 18:315–322.

Frost, I., W. P. Smith, S. Mitri, A. San Millan, Y. Davit, J. M. Osborne, J. M. Pitt-Francis, R. C. MacLean, and K. R. Foster. 2018. "Cooperation, competition and antibiotic resistance in bacterial colonies." The ISME journal 12:1582–1593.

Hamilton, W. D. 1964. "The genetical evolution of social behaviour. I." Journal of theoretical biology 7:1–16.

Harcombe, W. R., J. M. Chacón, E. M. Adamowicz, L. M. Chubiz, and C. J. Marx. 2018. "Evolution of bidirectional costly mutualism from byproduct consumption." Proceedings of the National Academy of Sciences 115:12000–12004.

Hart, S. F. M., D. Skelding, A. J. Waite, J. C. Burton, and W. Shou. 2019. "High-throughput quantification of microbial birth and death dynamics using fluorescence microscopy." Quantitative Biology 7:69–81.

Jansen, V. A., and M. Van Baalen. 2006. “Altruism through beard chromodynamics.” Nature 440:663-666.

Jayaraman, R. 2008. "Bacterial persistence: some new insights into an old phenomenon." Journal of biosciences 33:795–805.

Joshi, J., I. D. Couzin, S. A. Levin, and V. Guttal. 2017. "Mobility can promote the evolution of cooperation via emergent self-assortment dynamics." PLoS computational biology 13:e1005732.

Kim, J.-W. 2010. "A tag-based evolutionary prisoner’s dilemma game on networks with different topologies." Journal of Artificial Societies and Social Simulation 13:2.

Kim, J., and W. Park. 2015. "Indole: a signaling molecule or a mere metabolic byproduct that alters bacterial physiology at a high concentration?" Journal of Microbiology 53:421–428.

Kivisaar, M. 2020. "Mutation and recombination rates vary across bacterial chromosome." Microorganisms 8:25.

Laird, R. A. 2011. "Green-beard effect predicts the evolution of traitorousness in the two-tag Prisoner’s dilemma." Journal of theoretical biology 288:84–91.

Lee, H. H., M. N. Molla, C. R. Cantor, and J. J. Collins. 2010. "Bacterial charity work leads to population-wide resistance." Nature 467:82–85.

Lynch, M., M. S. Ackerman, J.-F. Gout, H. Long, W. Sung, W. K. Thomas, and P. L. Foster. 2016. "Genetic drift, selection and the evolution of the mutation rate." Nature Reviews Genetics 17:704–714.

Maynard-Smith, J. 1964. "Group Selection and Kin Selection." Nature 201:1145-1147.

Michod, R. E. 1982. "The theory of kin selection." Annual Review of Ecology and systematics 13:23-55.

Munita, J. M., and C. A. Arias. 2016. "Mechanisms of antibiotic resistance." Microbiology spectrum 4:4.2. 15.

Pepper, J. W., and B. B. Smuts. 2002. "A mechanism for the evolution of altruism among nonkin: positive assortment through environmental feedback." The American Naturalist 160:205–213.

Preussger, D., S. Giri, L. K. Muhsal, L. Oña, and C. Kost. 2020. "Reciprocal Fitness Feedbacks Promote the Evolution of Mutualistic Cooperation." Current Biology 30:3580–3590.e3587.

Sandoval-Motta, S., and M. Aldana. 2016. "Adaptive resistance to antibiotics in bacteria: a systems biology perspective." WIREs Systems Biology and Medicine 8:253–267.

Smith, P., and M. Schuster. 2019. "Public goods and cheating in microbes." Current Biology 29:R442–R447.

Thomas, C. M., and K. M. Nielsen. 2005. "Mechanisms of, and barriers to, horizontal gene transfer between bacteria." Nature Reviews Microbiology 3:711–721.

Trivers, R. L. 1971. "The Evolution of Reciprocal Altruism." The Quarterly review of biology 46:35–57.

Walter, T., K. H. Veldmann, S. Götker, T. Busche, C. Rückert, A. B. Kashkooli, J. Paulus, K. Cankar, and V. F. Wendisch. 2020. "Physiological response of Corynebacterium glutamicum to indole." Microorganisms 8:1945.

Wax, R. G., K. Lewis, A. A. Salyers, and H. Taber. 2007. Bacterial resistance to antimicrobials. CRC press.

West, S. A., G. A. Cooper, M. B. Ghoul, and A. S. Griffin. 2021. "Ten recent insights for our understanding of cooperation." Nature ecology & evolution 5:419–430.

West, S. A., A. S. Griffin, and A. Gardner. 2007. "Social semantics: altruism, cooperation, mutualism, strong reciprocity and group selection." Journal of evolutionary biology 20:415–432.

Wiuff, C., R. Zappala, R. Regoes, K. Garner, F. Baquero, and B. Levin. 2005. "Phenotypic tolerance: antibiotic enrichment of noninherited resistance in bacterial populations." Antimicrobial agents and chemotherapy 49:1483–1494.

Yanofsky, C., V. Horn, and P. Gollnick. 1991. "Physiological studies of tryptophan transport and tryptophanase operon induction in Escherichia coli." Journal of bacteriology 173:6009–6017.

Zarkan, A., J. Liu, M. Matuszewska, H. Gaimster, and D. K. Summers. 2020. "Local and universal action: the paradoxes of indole signalling in bacteria." Trends in microbiology 28:566–577.

