## Supplemental Figure 1 to 5 for "Phenotypic plasticity of antibiotic resistance, metabolism byproduct utilization and the evolution of mutually beneficial cooperation in *Escherichia coli*"

† authors contributed equally

1. **The cell number of *E. coli* in lysogeny broth (LB) solid medium with or without norfloxacin**

Wild-type *E.coli*-S was incubated for 12 hours in either 37.5, 75, 100 and 150 ng mL^-1^ (four replicates per treatment) norfloxacin culture medium in LB liquid medium. The OD_600_ of each replicate was measured and diluted to the same value. 50 µl of diluted culture was then inoculated to LB liquid medium with or without 600 ng mL^-1^. The OD_600_ value of each replicate was then measured and cell number calculated from OD_600_.

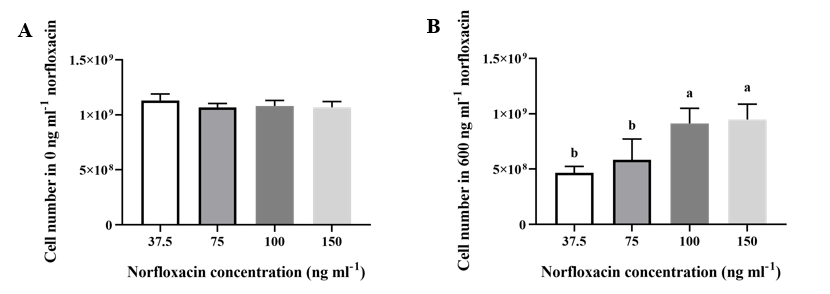

Figure S1. (A) After exposed to either 37.5, 75, 100, 150 ng mL^-1^ norfloxacin for 12 hours, the cell number of each culture in LB liuqid medium wihtout norfloxacin(incubation for 12 hours). (B) After exposed to either 37.5, 75, 100, 150 ng mL^-1^ norfloxacin for 12 hours, the cell number of each culture in LB medium with 600 ng mL^-1^ norfloxacin (12hours incubation). Results show means ± s.e. of four replicates per treatment, and different letters indicates significant difference (One-way anova, p = 0.002, F = 9.92, df = 15).

2. **Effect of different antibiotics on the ratio of antibiotic tolerant bacterial cells in LB liquid medium**

Wild-type *E. coli*-S was incubated for 12 hours at either a high or a low concentration of tetracycline, amplicillin or kanamycin in LB liquid medium. The proportion of resistant bacteria cells was measured by re-inoculation of each culture to LB medium with or without antibiotics (3 μg mL^-1^ tetracycline, 10 μg mL^-1^ ampicillin or 20 μg mL^-1^ kanamycin) for 12 hours. The ratio of resistant *E. coli* cells in the culture stressed with high level of antibiotics (tetracycline, amplicillin or kanamycin) was significantly higher (p<0.05, t-test) than in low concentration of antibiotics.

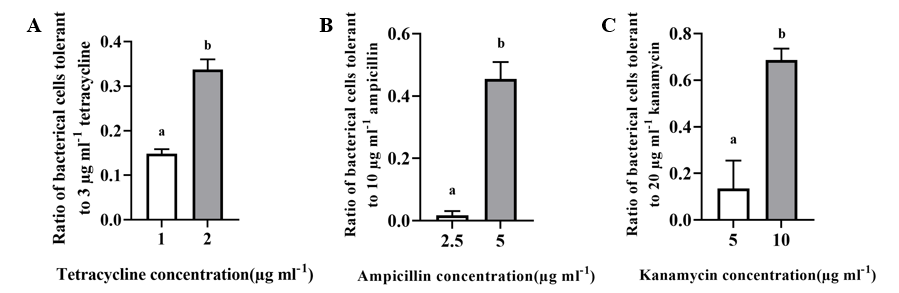

Figure S2. (A) The ratio of bacterial cells tolerant to 3 μg mL^-1^ tetracycline after 12 hours incubation in which samples were exposed for 12 hours to either 1, 2 μg mL^-1^ tetracycline. (B) the ratio of bacterial cells tolerant to 10 μg mL^-1^ ampicillin after 12 hours incubation in which the samples were exposed for 12 hours to either 2.5, 5 μg mL^-1^ ampicillin. (C) The ratio of bacterial cells tolerant to 20 μg mL^-1^ kanamycin after 12 hours incubation in which the samples were exposed to12 hours to either 5, 10 μg mL^-1^ kanamycin. Data presented are means ± 1 s.e. of three replicates per treatment, a and b indicates significant difference at p < 0.05 (t-test,tetracycline: t = -13.153,n = 3; Ampicillin: t = -13.39,n = 3; Kanamycin: t = -7.381,n = 3)

3. **Effect of different types antibiotics concentrations on the ratio of antibiotics tolerant bacterial cells in LB solid medium**

Wild-type *E. coli*-S was incubated for 12 hours at different concentrations of norfloxacin, tetracycline, amplicillin or kanamycin on LB liquid medium. The culture was then reinoculated to LB agar plates with or without a different type of antibiotics (3 μg mL^-1^ tetracycline, 600 ng mL^-1^ norfloxacin, 10 μg mL^-1^ ampicillin or 20 μg mL^-1^ kanamycin). The proportion of resistant cells in the total population on each agar plate was then measured by colony-forming unit (CFU) counting. The ratio of resistant cells in each culture stressed with a high level of antibiotics (tetracycline,norfloxacin, amplicillin or kanamycin ) was significantly higher (p<0.05, t-test) than for low concentrations of antibiotics.

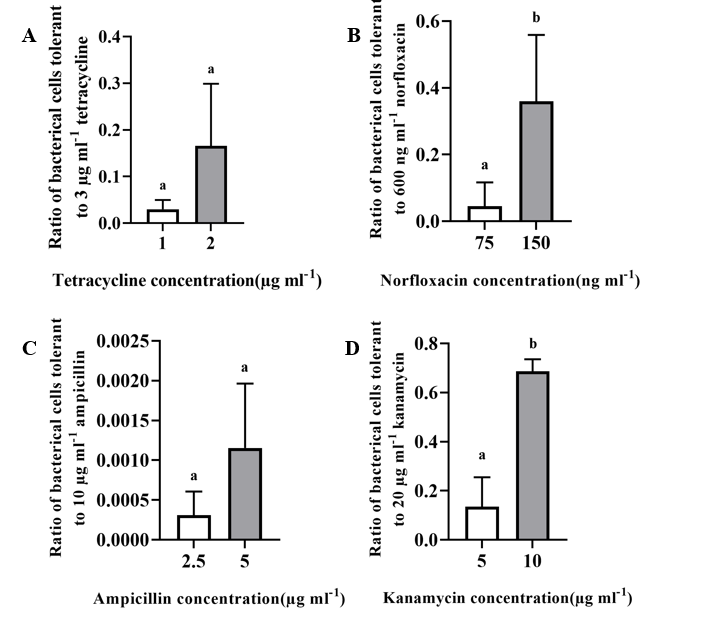

Figure S3. (A) The proportion of bacterial cells tolerant to 3 μg mL^-1^ tetracycline after exposed to either 1, 2 μg mL^-1^ Tetracycline for 12 hours. (B) proportion of bacterial cells tolerant to 600 ng mL^-1^ norfloxacin after exposed to 75, 150 ng mL^-1^ norfloxacin for 12 hours. (C) the proportion of bacterial cells tolerant to 10 μg mL^-1^ ampicillin after exposed to 2.5, 5 μg mL^-1^ ampicillin for 12 hours. (D) the proportion of bacterial cells tolerant to 20 μg mL^-1^ kanamycin after exposed to either 5 or 10 μg mL^-1^ kanamycin for 12 hours. Results were shown as average ± standard error of three replications, a and b indicates significant difference at p < 0.05 (Tetracycline:t= -1.714, n = 3; Norfloxacin: t = -3.309, n = 3; Ampicillin: t = -1.699, n = 3; Kanamycin: t = -7.138, n = 3).

4. **The effect of different concentrations of indole on the growth of *Escherichia coli* in lysogeny broth solid medium**

A different concentration of indole (0, 150, 300, 600 μ mol L^-1^) was added to the medium of *E. coli*-S. Growth (OD_600_) was measured after 24 hours incubation. There was no significant difference on growth attributable to variation in indole (Fig S4; Kruskal-Wallis test: p = 0.502, H = 2.356, n = 12)

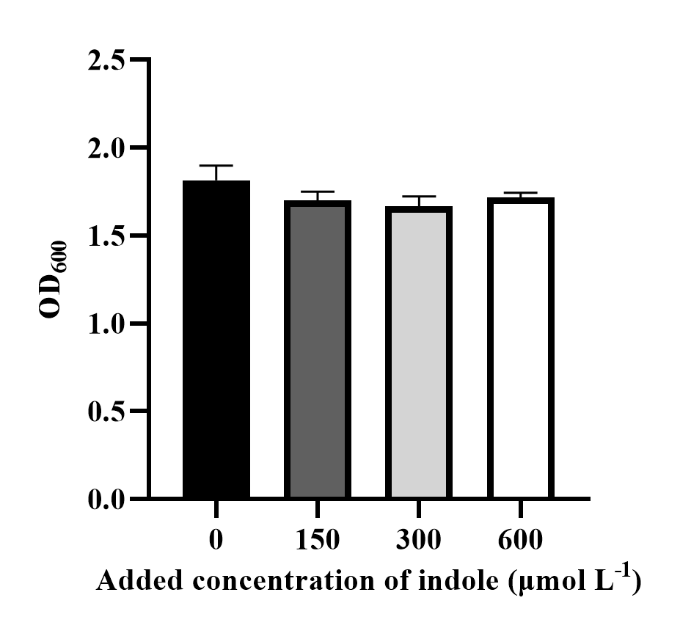

Figure S4. Growth of *E. coli*-S measured by OD_600_ after 24 hours culturing in LB liquid medium at different concentrations of indole (0, 150, 300 and 600 μ mol L^-1^). Results were shown as average ± standard error of three replications of four treatments.

The effect of different concentrations of indole (0, 150, 300, 600 μ mol L^-1^) on growth of *E. coli*-R with or without the presence of norfloxacin after 24 hours incubation was also measured (Figure S5). When incubated with 2400 ng mL^-1^ norfloxacin, both 300 and 600 μmol L^-1^ indole addition significantly inhibited *E. coli-*R growth from 1.09±0.11 to 0.94±0.04 (One way anova, p = 0.041, n = 3) and 0.86±0.04 (one way anova, p = 0.004, n = 3), respectively (Figure S5A). When *E. coli-*R was incubated without norfloxacin, the growth of *E. coli-*R was significantly (One way anova, p = 0.041, n = 3) inhibited from 1.15±0.16 in control treatment to 0.93±0.02 in 600 μmol L^-1^ indole treatment (Figure S5B).

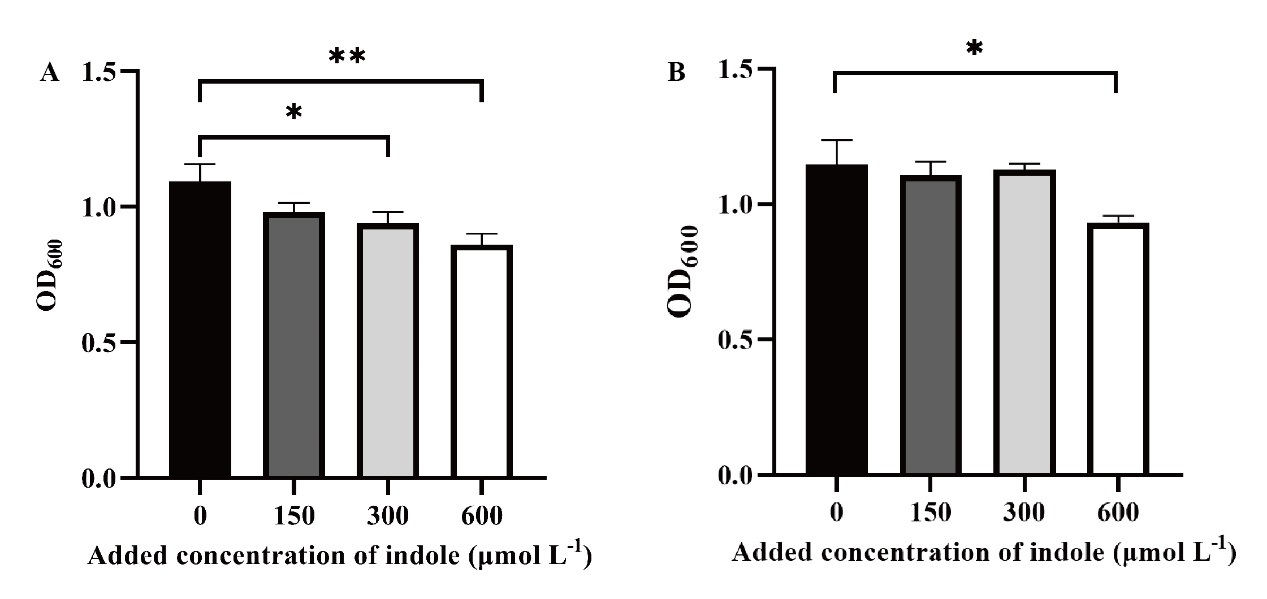

Figure S5. The effect of variation in exposure to indole (0, 150, 300 and 600 μ mol L^-1^) on growth of *E. coli-R* measured by OD_600_ after 24 hours culturing in LB liquid medium with (A) or without (B) 2400 ng mL^-1^ norfloxacin. Results were shown as average ± standard error of three replications of four treatments. ** indicates significant difference at p < 0.01 and * indicates significant difference at p < 0.05.
